# The molecular architecture of cell cycle arrest

**DOI:** 10.1101/2022.04.27.489684

**Authors:** Wayne Stallaert, Sovanny R. Taylor, Katarzyna M. Kedziora, Martha S. Johnson, Colin D. Taylor, Holly K. Sobon, Catherine L. Young, Juanita C. Limas, Jonah Varblow Holloway, Jeanette Gowen Cook, Jeremy E. Purvis

**Affiliations:** Department of Genetics, University of North Carolina at Chapel Hill, Chapel Hill, NC, USA; Computational Medicine Program, University of North Carolina at Chapel Hill, Chapel Hill, NC, USA; Bioinformatics and Analytics Research Collaborative (BARC), University of North Carolina at Chapel Hill, Chapel Hill, NC, USA; Department of Biochemistry and Biophysics, University of North Carolina at Chapel Hill, NC, USA; Department of Pharmacology, University of North Carolina at Chapel Hill, Chapel Hill, NC, USA

**Author notes:** Corresponding Author: Jeremy Purvis Mary Ellen Jones Building 11018C, CB#7488 116 Manning Drive Chapel Hill, NC 27599-7488.

## Abstract

The cellular decision governing the transition between proliferative and arrested states is crucial to the development and function of every tissue. While the molecular mechanisms that regulate the proliferative cell cycle are well established, we know comparatively little about what happens to cells as they diverge into cell cycle arrest. We performed hyperplexed imaging of 49 cell cycle effectors to obtain a map of the molecular architecture that governs cell cycle exit and progression into reversible (“quiescent”) and irreversible (“senescent”) arrest states. Using this map, we found multiple points of divergence from the proliferative cell cycle; identified stress-specific states of arrest; and resolved the molecular mechanisms governing these fate decisions, which we validated by single-cell, time-lapse imaging. Notably, we found that cells can exit into senescence from either G1 or G2; however, both subpopulations converge onto a single senescent state with a G1-like molecular signature. Cells can escape from this “irreversible” arrest state through the upregulation of G1 cyclins. This comprehensive map provides a first glimpse of the overall organization of cell proliferation and arrest.

## Introduction

The decision of when and where to trigger cell division is fundamental to nearly all aspects of development and physiology. At the level of the individual cell, the molecular basis of the proliferation/arrest decision is embedded within a highly interconnected and dynamic network of cell cycle regulators. Progression through the proliferative phases of the cell cycle (G1/S/G2/M) is governed by a series of biochemical reactions that are coordinated in time and space to ensure the successful replication of DNA and its division into two daughter cells. In addition to these four proliferative phases, cells may also “exit” the proliferative cell cycle into a state of cell cycle arrest, often referred to as G0. While arrested, cells still perform many essential cellular functions including metabolism, secretion, transcription, and translation. However, as long as they remain in the G0 state, arrested cells neither synthesize DNA nor undergo cell division. This five-state model has become the canonical cell cycle model found in most textbooks (Morgan, 2007) and the current literature (Spencer *et al*, 2013; Overton *et al*, 2014; Marescal & Cheeseman, 2020) and has shaped our thinking about the cell cycle for over 70 years (Howard & Pelc, 1951; Cameron & Greulich, 1963; Smith & Martin, 1973).

While the mechanisms that govern progression through the proliferative cell cycle have been studied extensively, we know comparatively little about what happens to cells after they exit the proliferative cell cycle. We know that cells may exit the cell cycle in response to various biochemical (e.g. DNA damage, oxidative stress) or environmental insults (e.g. lack of mitogens, high local cell density) triggered by different molecular mechanisms (Sagot & Laporte, 2019; Marescal & Cheeseman, 2020). After exiting the cell cycle, cells may progress into deeper states of reversible (“quiescent”) cell cycle arrest (Owen *et al*, 1989; Wang *et al*, 2017; Kwon *et al*, 2017), and in some cases can transition into an irreversible (“senescent”) state of arrest (Sousa-Victor *et al*, 2014; Marthandan *et al*, 2014; Fujimaki *et al*, 2019; Fujimaki & Yao, 2020). Clearly, cell cycle arrest is far from a single, static molecular state (Klosinska *et al*, 2011; Sun & Buttitta, 2017; Coller *et al*, 2006), yet a systematic characterization of when and how cells arrest remains lacking.

In this study, we used a combination of hyperplexed, single-cell imaging and manifold learning to map the molecular architecture of cell cycle arrest. Previously, we used this approach to map the structure of the proliferative cell cycle in unperturbed, non-transformed retinal pigment epithelial (RPE) cells (Stallaert *et al*, 2021). Building on this work, we exposed asynchronous RPE cells to three distinct stressors—hypomitogenic, replication and oxidative—known to induce cell cycle arrest. For each stress, we identify the points of exit from the proliferative cell cycle, the mechanism that induced arrest, and the molecular signatures of cells as they transition through distinct arrest states. We reveal that cells exit the cell cycle along two distinct arrest trajectories in response to replicative and oxidative stress and that these trajectories are distinct from the arrest state induced by hypomitogenic stress. We show how sustained replication stress generates polyploidy through mitotic skipping and endoreduplication. Finally, we identify the molecular trajectories that lead to “irreversible” arrest and reveal that cellular senescence is an obligate G1-like molecular state that can be reversed by increasing the expression of G1 cyclins.

## Results

To map the molecular architecture of cell cycle arrest, we subjected an asynchronous population of RPE cells to a variety of natural stresses known to induce exit from the proliferative cell cycle. These stresses included hypomitogenic stress (induced by serum starvation), replication stress (using the topoisomerase inhibitor etoposide) and oxidative stress (by exogenous H_2_O_2_ addition). We performed iterative indirect immunofluorescence imaging (4i) (Gut *et al*, 2018) of 49 cell cycle effectors (**Table EV1**) and extracted 2952 unique single-cell features from this imaging dataset. These features included the subcellular expression of each protein across different cellular compartments (i.e. nucleus, cytosol, plasma membrane and perinuclear region) as well as cell morphological features, such as size and shape, for 23,605 individual cells (**Fig 1A**). After feature selection (to eliminate features that vary in a cell-cycle-independent manner (Stallaert *et al*, 2021)), we performed manifold learning using Potential of Heat-diffusion for Affinity-based Transition Embedding (PHATE) (Moon *et al*, 2019) to find the “surface” within this high-dimensional space that represents progression through cell cycle space and visualize it as a 2-dimensional representation. Using these lower dimensional representations or cell cycle “maps”, we were able to identify the points at which cells exit the proliferative cell cycle in response to each stress and the mechanisms governing these proliferation/arrest decisions.

**Fig 1:**
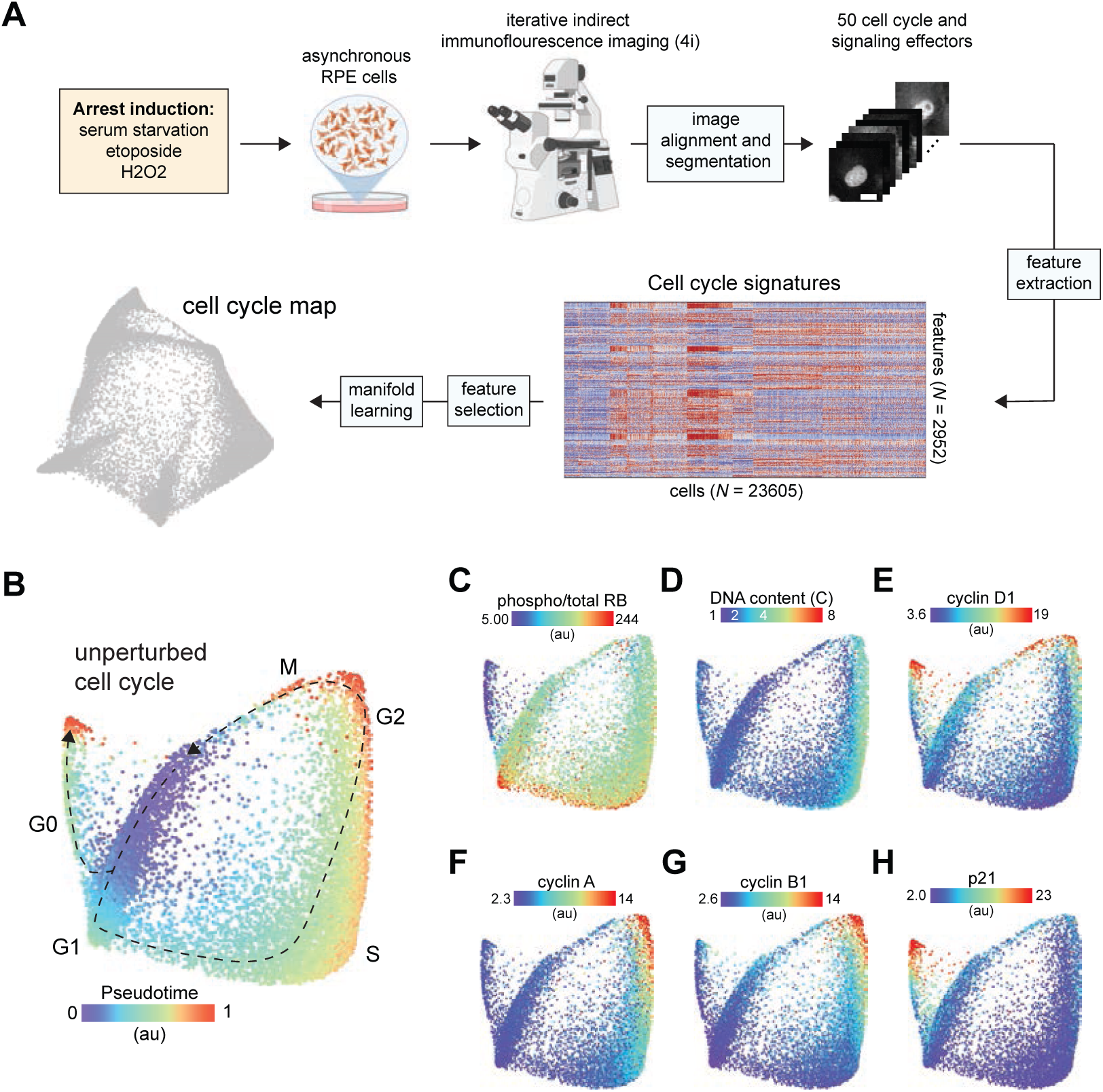
Mapping the architecture of cell cycle arrest. A. Schematic of the experimental approach. B. Cell cycle map of unperturbed cells (*N* = 11 268 cells). Diffusion pseudotime values are plotted and inferred proliferative (G1,S,G2,M) and arrest trajectories (G0) are indicated on the map. C-H. (C) Phospho/total RB, (D) DNA content, (E) cyclin D1, (F) cyclin A, (G) cyclin B1 and (H) p21 of unperturbed cells are plotted on the map. Median nuclear values are shown

To identify the precise molecular states in which proliferating cells exit the cell cycle, we first resolved the unperturbed cell cycle as a reference map. In unperturbed RPE cells, we obtained a map that consisted of a cyclical proliferative trajectory (phospho-RB-positive) that proceeds sequentially through the canonical phases of the cell cycle (G1/S/G2/M) and an arrest trajectory (phospho-RB-negative) that diverges from the proliferative trajectory soon after cell division (**Fig 1B-G**), which is consistent with the cell cycle map obtained in a previous study (Stallaert *et al*, 2021) and was reproducible across experimental replicates (**Fig EV1**). Previous studies indicate that cells may enter a “spontaneous” state of cell cycle arrest shortly after cell division due to low levels of endogenous stress (including replication stress) during the mother cell cycle (Arora *et al*, 2017; Min & Spencer, 2019), driven by an increase in p21 expression (**Fig 1H**).

### Hypomitogenic stress

To induce hypomitogenic stress, cells were serum starved for 1 or 7 days prior to fixation. To show how hypomitogenic stress disrupts the normal cycling of cells, we generated a unified cell cycle map of both unperturbed (**Fig 2A, dark gray**) and serum-starved cells (**Fig 2A, light and dark blue**). This new embedding shows the proliferative and arrest trajectories of unperturbed cells (from **Fig 1**) and indicates that the state of cell cycle arrest induced by serum starvation is distinct from the spontaneous arrest observed in unperturbed cells. Trajectory inference by diffusion pseudotime revealed that serum-starved cells diverge from the proliferative cell cycle during G2 (**Fig 2B**). However, unlike spontaneous arrest, this cell cycle exit was not accompanied by a large increase in p21 (**Fig 2C**). In the unperturbed cell cycle, cyclin D1 increased in late G2 and remained elevated during mitosis and after cell division (**Fig 2D, left panel**) (Gookin *et al*, 2017). In the presence of hypomitogenic stress, however, cyclin D1 remained low during G2 (**Fig 2D, right panel**) and cells underwent mitosis (as indicated by a drop in DNA content) (**Fig 2E**) directly into a state of arrest (**Fig 2F**). To validate this mechanism of cell cycle exit, we performed time-lapse imaging of RPE cells expressing endogenous cyclin D1 tagged with a fluorophore at its endogenous locus (cycD1-Venus), as well as a fluorescent biosensor of CDK2 activity (DHB-mCherry), which can be used to distinguish actively proliferative versus arrested cells (Spencer *et al*, 2013). While unperturbed cells exhibited a clear increase in cyclin D1 during G2, serum starvation significantly reduced the induction of cyclin D1 during G2, leading to less inherited cyclin D1 in daughters cells (**Fig 2G**) (Guo *et al*, 2005).

**Fig 2:**
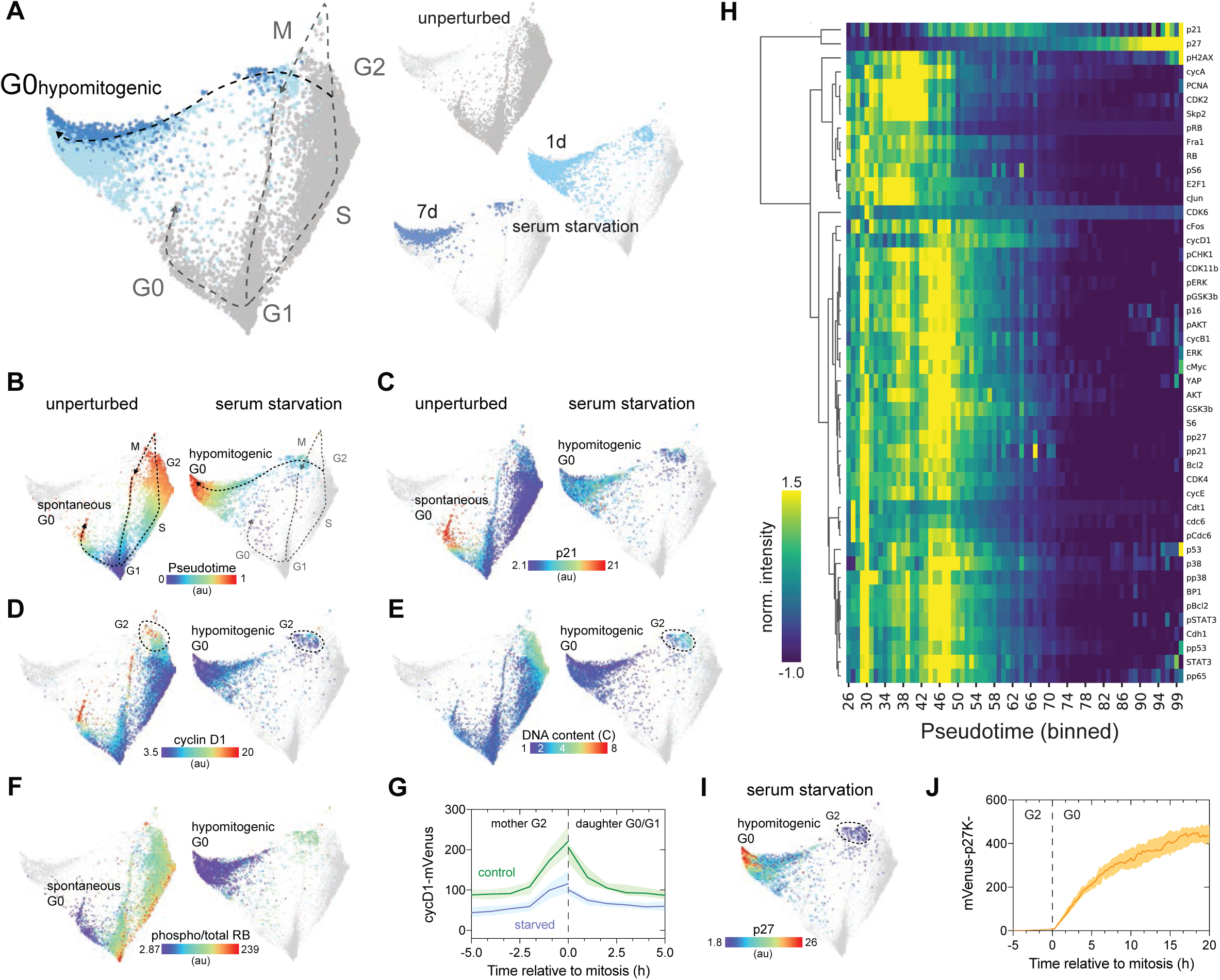
The arrest architecture of hypomitogenic stress. A. Unified cell cycle map of unperturbed (gray) and serum-starved cells (1d: light blue, 7d: dark blue, *N* = 3007 cells). The proliferative cell cycle (dotted gray line) and the hypomitogenic arrest trajectory (black dotted line) are indicated on the map. Inset: Each treatment condition is shown individually on the unified map (other conditions are shown in lighter gray). B-F. (B) Diffusion pseudotime, (C) p21, (D) cyclin D1, (E) DNA content and (F) phospho/total RB of unperturbed (left panels) or serum-starved cells (right panels) are plotted on the map. G. Time-lapse imaging of cyclinD1-mVenus intensity in unperturbed (control) and serum-starved cells. *N =* 105 control cells and *N* = 111 starved cells. H. Heatmap of feature intensity along the hypomitogenic arrest trajectory. Features were ordered by hierarchical clustering according to their dynamics along the arrest trajectory. Diffusion pseudotime values were binned and pseudotime values with <15 cells were excluded from the visualization. I. Median nuclear p27 abundance in serum-starved cells is plotted on the map. J. Time-lapse imaging of mVenus-p27K-intensity in serum-starved cells. *N* = 101 cells. For time-lapse imaging data, the population median (thick line) +/-95% confidence interval (shaded areas) are shown.

After exiting the cell cycle, progression further along the hypomitogenic arrest trajectory was accompanied by a considerable decrease in the abundance of nearly every protein measured including key proliferative effectors such as CDK2, CDK4, CDK6, CDH1, CDT1, PCNA, SKP2, FRA1 and cJUN, as well as decreased nuclear YAP and mTOR signaling (S6 phosphorylation) (**Fig 2H**). The only proteins not downregulated following serum starvation were the CDK inhibitor proteins p27 (**Fig 2H-I**) and, to a much lesser extent, p21 (**Fig 2C** **and** **Fig 2H**). In fact, the abundance of p27 increased as cells progressed further along the arrest trajectory. Using a fluorescent biosensor for p27 (mVenus-p27K-) (Oki *et al*, 2014), we confirmed that p27 increased steadily in individual cells following serum starvation, starting immediately after cell division (**Fig 2J**) (Coats *et al*, 1996).

### Replication stress

To induce replication stress, cells were treated with etoposide (1 μM), an inhibitor of DNA topoisomerase II that interferes with DNA re-ligation step during replication, for 1-4 days prior to fixation. Once again, we constructed a unified cell cycle map of unperturbed (**Fig 3A, dark gray**) and etoposide-treated cells (**Fig 3A, green**) to show how replicative stress interferes with cell cycle progression. Within a single population of cells treated with etoposide, individual cells diverged from the proliferative cell cycle along two distinct arrest trajectories. One subpopulation exited from a G2-like state after DNA replication was complete (DNA content = 4C). A second subpopulation exited the cell cycle in the subsequent G1 phase of daughter cells immediately following mitosis (DNA content = 2C) (**Fig 3B-C**). Both subpopulations entered arrest states characterized by a loss of RB phosphorylation (**Fig 3D**). Cell cycle exit along the 4C trajectory was accompanied by activation of the DNA damage checkpoint in G2 as indicated by an increase in markers of DNA damage signaling, including phospho-H2AX, phospho-CHK1, phospho-p65, p53 (**Fig EV2A-E**) and p21 (**Fig 3E**). In contrast, daughter cells that exited the cell cycle following mitosis along the 2C trajectory did not express early markers of DNA damage signaling (phospho-H2AX, phospho-CHK1) (**Fig EV2B-C**), but possessed sustained elevation of phospho-p65, p53 (**Fig EV2D-E**) and p21 (**Fig 3E**), consistent with replication stress inherited from the previous cell cycle (Arora *et al*, 2017).

**Fig 3:**
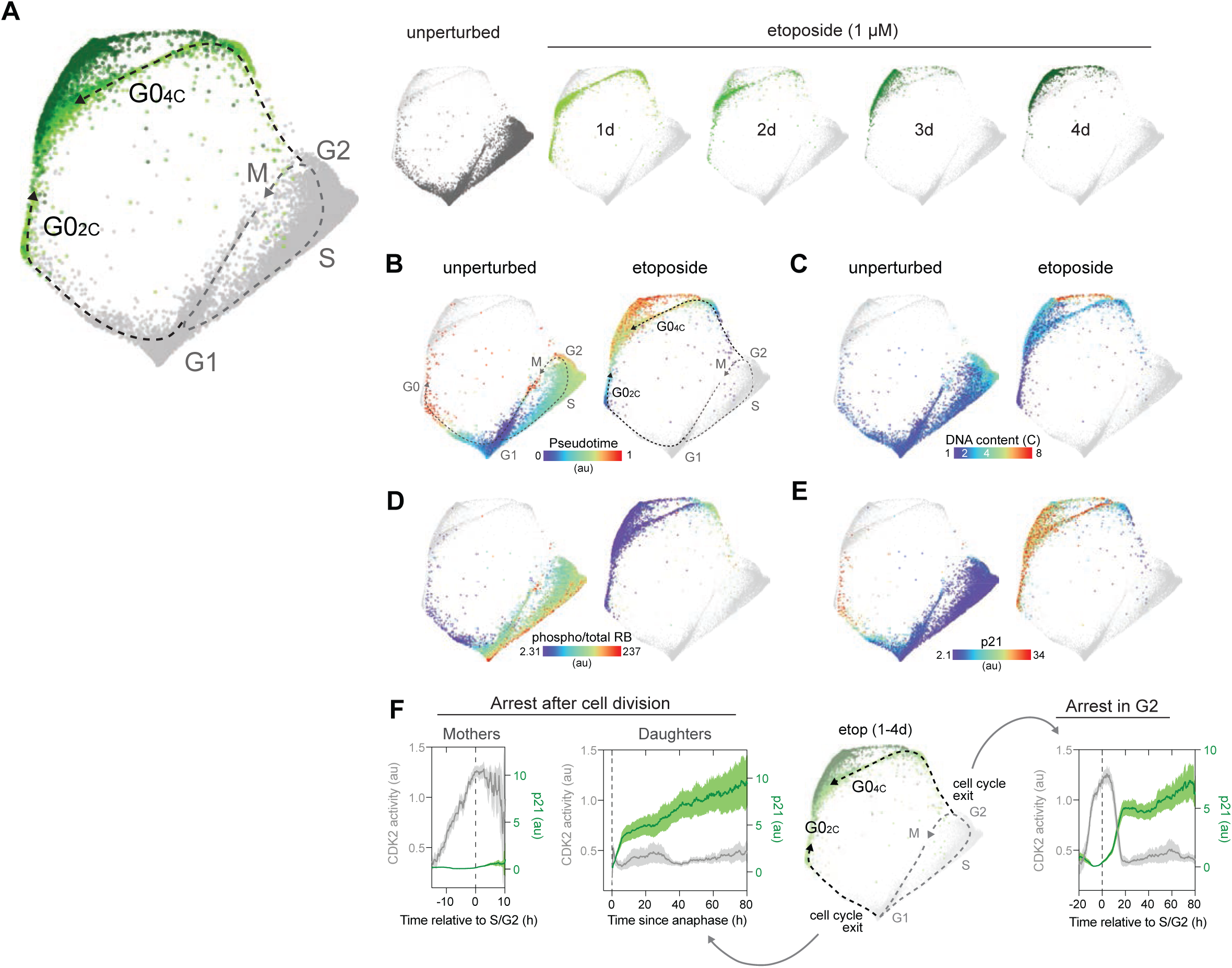
The arrest architecture of replication stress. A. Unified cell cycle map of unperturbed (gray) and etoposide-treated cells (1 μM; 1d: light green, 2d: green, 3d: dark green, 4d: darker green – see inset, *N* = 4315 cells). The unperturbed cell cycle (dotted gray line) and two arrest trajectories (into G0_2C_ and G0_4C_; black dotted lines) are indicated on the map. Inset: Each condition is shown individually on the map (other conditions are shown in lighter gray). B-E. (B) Diffusion pseudotime, (C) DNA content, (D) phospho/total RB and (E) p21 of unperturbed (left panels) or etoposide-treated cells (right panels) are plotted on the arrest architecture. Median nuclear values are shown. F. Time-lapse imaging of CDK2 activity (DHB-mCherry, gray) and p21-YPet (green) intensity in etoposide-treated cells. Schematic shows the two arrest trajectories observed following etoposide treatment. Cells that successfully complete G2 (“Mothers”, *N* = 32 cells) but arrest following cell division (“Daughters”, *N* = 45 cells) are shown in the two leftmost panels. Cells that arrest in G2 (*N =* 40 cells) are shown in the rightmost panel. Population median (thick line) and individual traces (thin lines) are shown.

To validate the observation that individual cells exit the cell cycle along two distinct arrest trajectories in response to replication stress and to investigate the mechanisms that govern this decision, we performed single cell time-lapse imaging of RPE cells expressing a cell cycle sensor (PCNA-mTq2), a CDK2 activity sensor to detect cell cycle arrest (DHB-mCherry), and endogenous p21 fused to a fluorophore (p21-YPet) for 4 days after etoposide treatment. We observed a similar bifurcation of cell fate in live cells in response to replication stress, with 56% of cells exiting the cell cycle in G2 (along the 4C trajectory) and 44% proceeding through to mitosis following etoposide treatment (**Fig 3F**). The loss of CDK2 activity that accompanied cell cycle arrest in G2 occurred simultaneously with an increase in p21 expression (**Fig 3F, last panel**). For each of the cells that successfully progressed through to mitosis, however, no detectable p21 induction was observed in G2 (**Fig 3F, first panel**). Instead, their daughter cells arrested immediately following cell division (as indicated by a sustained decrease in CDK2 activity) accompanied by an increase in p21 expression shortly after cell division (**Fig 3F, second panel**). Over several days of etoposide treatment cells proceed further along the 2C and 4C arrest trajectories, transitioning through additional molecular states (**Fig 3A** **inset**), which we will discuss in greater detail below.

### Oxidative stress

To assess how the cell cycle responds to oxidative stress, cells were treated with hydrogen peroxide (H_2_O_2_) for 1, 2 or 3 days. Overall, the arrest architecture of oxidative stress was very similar to that induced by replication stress (**Fig EV3A**), suggesting that the dominant cell cycle response to exogenous oxidative stress is primarily related to its ability to induce DNA damage (Demple & Halbrook, 1983). Consistent with this notion, oxidative stress induced elevated markers of the DNA damage response including phospho-H2AX, phospho-CHK1, p53 and p21 (**Fig EV3B-E**). Similar to etoposide treatment, H_2_O_2_ induced cell cycle exit along two distinct trajectories diverging from either G1 or G2, into arrest states (**Fig EV3F**) with 2C or 4C DNA content (**Fig EV3G**), respectively, both accompanied by p21 induction (**Fig EV3E**). However, unlike etoposide, H_2_O_2_ is rapidly metabolized following addition to cells (Sobotta *et al*, 2013) and thus the resultant DNA damage is more transient. As a result, markers of DNA damage decreased more rapidly after treatment (**Fig EV3H**) and a higher proportion of cells remained in the cell cycle over time (**Fig EV3I**) compared to the sustained replication stress induced by etoposide.

### Senescence, mitotic skipping and polyploidy

Over the 4 days of etoposide treatment, cells transitioned through different arrest states (**Fig 4A-B**) accompanied by changes in their molecular signatures (**Fig EV2F-G**). In the first 1-2 days, cells populated multiple, distinct arrest trajectories (“2C state” and “4C state”) (**Fig 4B**). Over time, however, most cells transitioned further along these trajectories and eventually converged on a single region of the structure (“terminal state”), while a small proportion of cells also began to populate a region consisting entirely of polyploidy cells (“8C state”). Similarly, after ∼3-4 days of etoposide treatment, most cells begin to possess elevated senescence-associated β-galactosidase (SA-β-gal) activity (**Fig 4C**), a hallmark of senescence (Hjelmeland *et al*, 1999). We therefore hypothesized that these states may represent irreversibly arrested senescent states. We previously identified a multivariate, proteomic signature of senescence in RPE cells (Stallaert *et al*, 2021). We used this signature to identify senescent cells following etoposide treatment. Cells in the terminal state possessed many of these senescent markers including GSK3β, phospho(Thr157)- and total p27, CDK4, cyclin D1 and cyclin E (**Fig 4D-I**). This subpopulation contained the largest cells in the population (**Fig 4J**) and possessed the lowest DNA:cytoplasm ratio (**Fig 4K**), another hallmark of senescent cells (Neurohr *et al*, 2019). Cells in the 8C state expressed some of these senescent features but were notably lacking p27 and CDK4, suggesting that these arrested polyploid cells reside in a different molecular state than the senescent cells in the terminal region.

**Fig 4:**
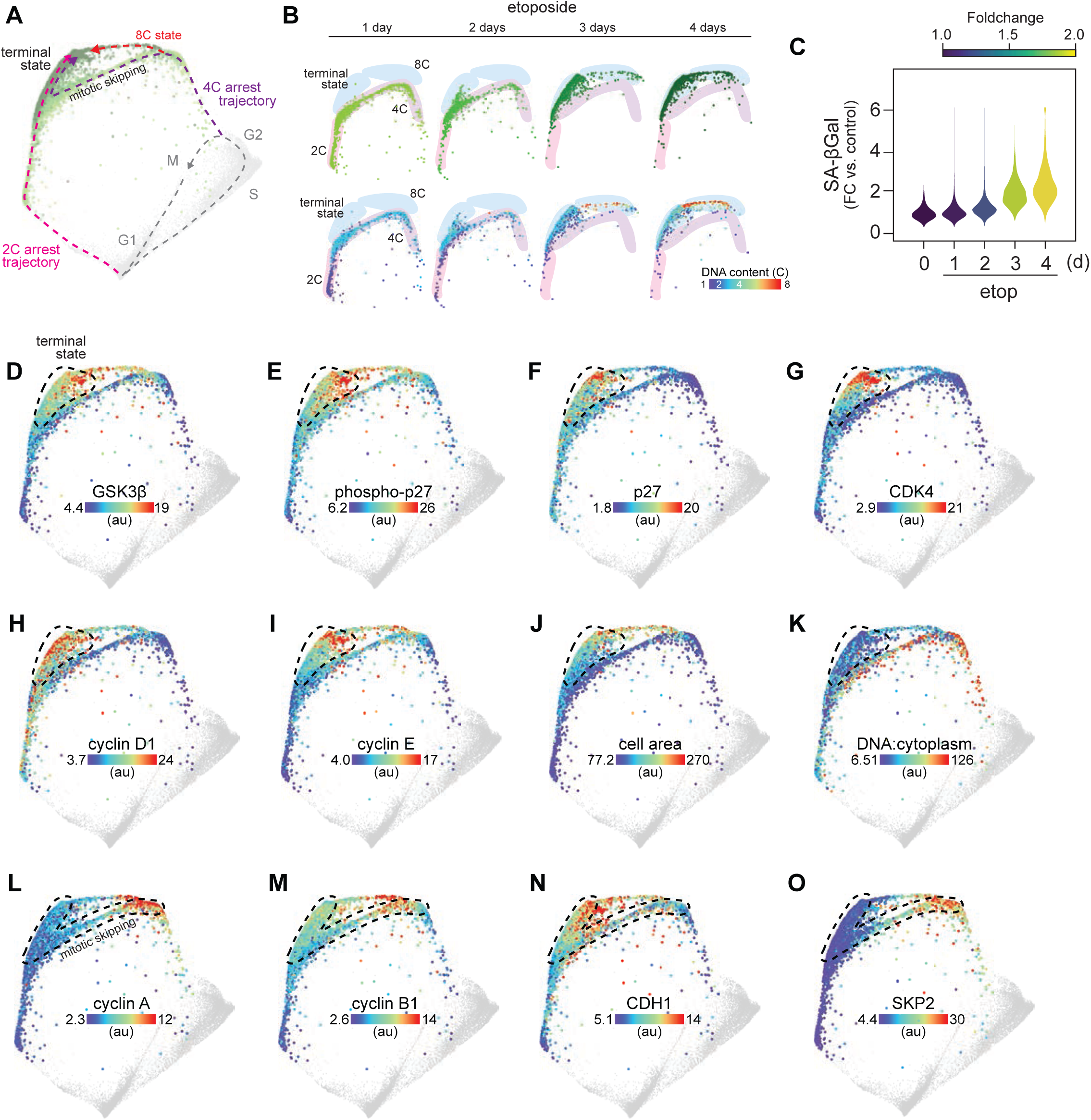
Replication stress promotes mitotic skipping and cellular senescence. A. Arrest architecture of replicative stress. 2C and 4C arrest trajectories are shown, the regions that correspond to mitotic skipping and the senescent state are annotated and the arrest trajectory of polyploid cells is indicated. B. Progression of arrest states over time following etoposide treatment. Etoposide-treated cells (upper panel: colored by condition, lower panel: colored by DNA content) populating 2C, 4C, 8C and senescent arrest states at each day of treatment (1-4 d). C. Distribution of senescence-associated β-galactosidase (SA-βgal) activity in individual cells following etoposide treatment (1 μM, 0-4 d). D-K. (D) GSK3β, (E) phospho(Thr157)-p27, (F) p27, (G) CDK4, (H) cyclin D1, (I) cyclin E, (J) cell area and (K) DNA:cytoplasm ratio of etoposide-treated cells are plotted on the map. Median nuclear values are shown in D-I. Area indicated with a dotted line represents the senescent region of the map. L-O. (L) Cyclin A, (M) cyclin B1, (N) CDH1 and (O) SKP2 of etoposide-treated cells are plotted on the map. Median nuclear values are shown. Area indicated with a dotted line shows the trajectory of mitotic skipping and transition into senescence.

We next investigated how cells that exited the cell cycle from G2 (with 4 copies of DNA) converged on a molecular state expressing high levels of G1 cyclins and CDKs. As cells progressed along the G2 arrest trajectory, there was an abrupt degradation of the G2/M cyclins A and B (**Fig 4L-M**) that coincided with the loss of RB phosphorylation (**Fig 3D**). Degradation of G2/M cyclins normally occurs during mitosis (Glotzer *et al*, 1991); however, we observed no change in DNA content (**Fig 3C**) nor any visual evidence of mitotic events in the images of cells along this trajectory. Progression along the G2 trajectory was also accompanied by an increase in APC/C subunit CDH1 and a loss of SKP2 (**Fig N-O**). These molecular events are consistent with a transition into a pseudo-G1 molecular state through a phenomenon known as “mitotic skipping” (**Fig 4A**), which often precedes the transition into senescence (Suzuki *et al*, 2012; Johmura *et al*, 2014). This phenomenon did not appear to be specific to RPE cells, as we observed that etoposide treatment induced a pseudo-G1 senescent state (high cyclin D1, low cyclin A and 4 copies of DNA) in osteosarcoma (U-2 OS) cells (**Fig EV4**) as well.

Next, we asked how the molecular signature of this pseudo-G1 senescent state compares to a *bone fide* G1 senescent state. Acute CDK4/6 inhibition with palbociclib (1 μM, 24 h) triggered cell cycle exit from G1 into a state of arrest distinct from hypomitogenic, replication or oxidative stress (**Fig EV5A**). Sustained palbociclib treatment (4-8 d) induced senescence as measured by an increase in SA-β-gal activity (**Fig 5A**). To compare the G1 and pseudo-G1 senescent states induced by palbociclib and etoposide, respectively, we performed a targeted 4i experiment making single-cell measurements of 37 cell cycle and signaling effectors (including cell cycle features previously shown to be upregulated in senescent cells (Stallaert *et al*, 2021)), as well as several measures of the senescence-associated secretory phenotype (SASP). We then used manifold learning to map the trajectories of cell cycle exit and progression into each senescent state (**Fig 5B**). Again, we observed that etoposide induced cell cycle exit from both G1 and G2, while palbociclib induced cell cycle exit from G1 along a distinct arrest trajectory (**Fig 5B-D**). These arrest trajectories, however, all converge on a single terminal state possessing elevated expression of the CDK inhibitors p16 and p27, SASP factors TGFβ1, IL-6 and IL-8, as well as activation of NF-κb (phospho-p65), AKT, p38, BCL-2, STAT3, STAT5 and SMAD2 (**Fig 5E-K, Fig EV5B-F**).

**Fig 5:**
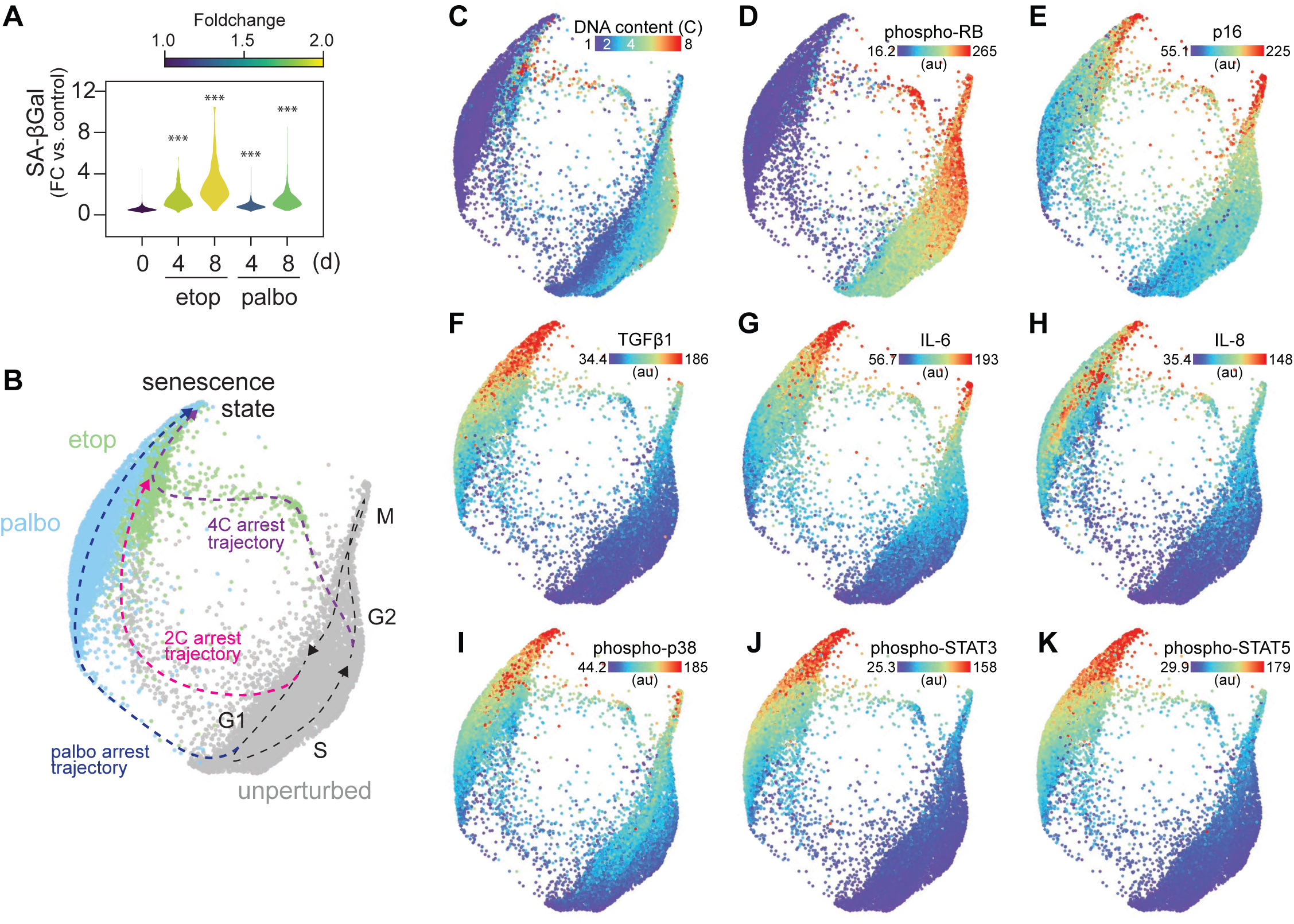
Palbociclib- and etoposide-induced arrest trajectories converge on a single state of cellular senescence. A. Unified cell cycle map of the unperturbed (gray), palbociclib-(palbo, 1 μM, 4/8 d) and etoposide- (etop,1 μM, 4/8 d) treated cells. The proliferative cell cycle trajectory from unperturbed cells was annotated using changes in DNA content (panel C) and cyclin abundance (not shown). Control: *N* = 10499 cells, etop: *N* = 2692 cells, palbo: *N* = 4931 cells B. Distribution of senescence-associated β-galactosidase (SA-βgal) activity in individual cells following etoposide or palbociclib treatment (1 μM, 4/8 d). Control: *N* = 6050 cells, etop 4d: *N* = 409 cells, etop 8d: *N* = 732 cells, palbo 4d: *N* = 1837 cells, palbo 8d: *N* = 1677 cells. Statistical significance was determined using using a one-way analysis of variance (ANOVA) with Sidak’s post hoc test (*** *p* < 0.0001). C-K. (C) DNA content, (D) phospho/total RB, (E) p16, (F) TGFβ1, (G) IL-6, (H) IL-8, (I) phospho-p38, (J) phospho-STAT3 and (K) phospho-STAT5 are plotted on the map. Median nuclear values are shown.

Our results suggest that the cell cycle status of senescent cells is a G1-like state. If true, then the mechanisms that maintain this arrested state, and in turn govern the molecular routes to possible cell cycle re-entry, should be limited to G1 regulatory events. Consistent with this hypothesis, we found that the upward trajectory through the senescent region of the replication stress arrest architecture (**Fig 6A**) showed molecular changes consistent with cell cycle re-entry in G1 and progress toward S phase including sequential increases in cyclin D1 (**Fig 4H**), cyclin E (**Fig 4I**), the DNA licensing factor Cdt1 (**Fig 6B**), and E2F activity (**Fig 6C**). As previously mentioned, we also observed a gradual accumulation of polyploid cells with 8C DNA content after 3-4 days of etoposide treatment (**Fig 4A-B**, **Fig 6A**). We therefore hypothesized that cells might be able to re-enter the cell cycle from the terminal state following mitotic skipping and undergo a second round of DNA replication, or “endoreduplication” (Fox & Duronio, 2013). To validate these observations, we performed time-lapse imaging of RPE cells expressing p21-YPet as well as cell cycle (PCNA-mTq2) and CDK2 activity (DHB-mCherry) sensors for 4 days following etoposide treatment. We indeed observed cells exiting the cell cycle from G2, remaining arrested for an extended period of time (15-30h), then re-entering the cell cycle (as indicated by an increase in CDK2 activity) and transitioning into a second S phase (as indicated by an increase in PCNA foci) without first undergoing mitosis (5/117 cells over 4 days of imaging; **Fig 6D**).

**Fig 6:**
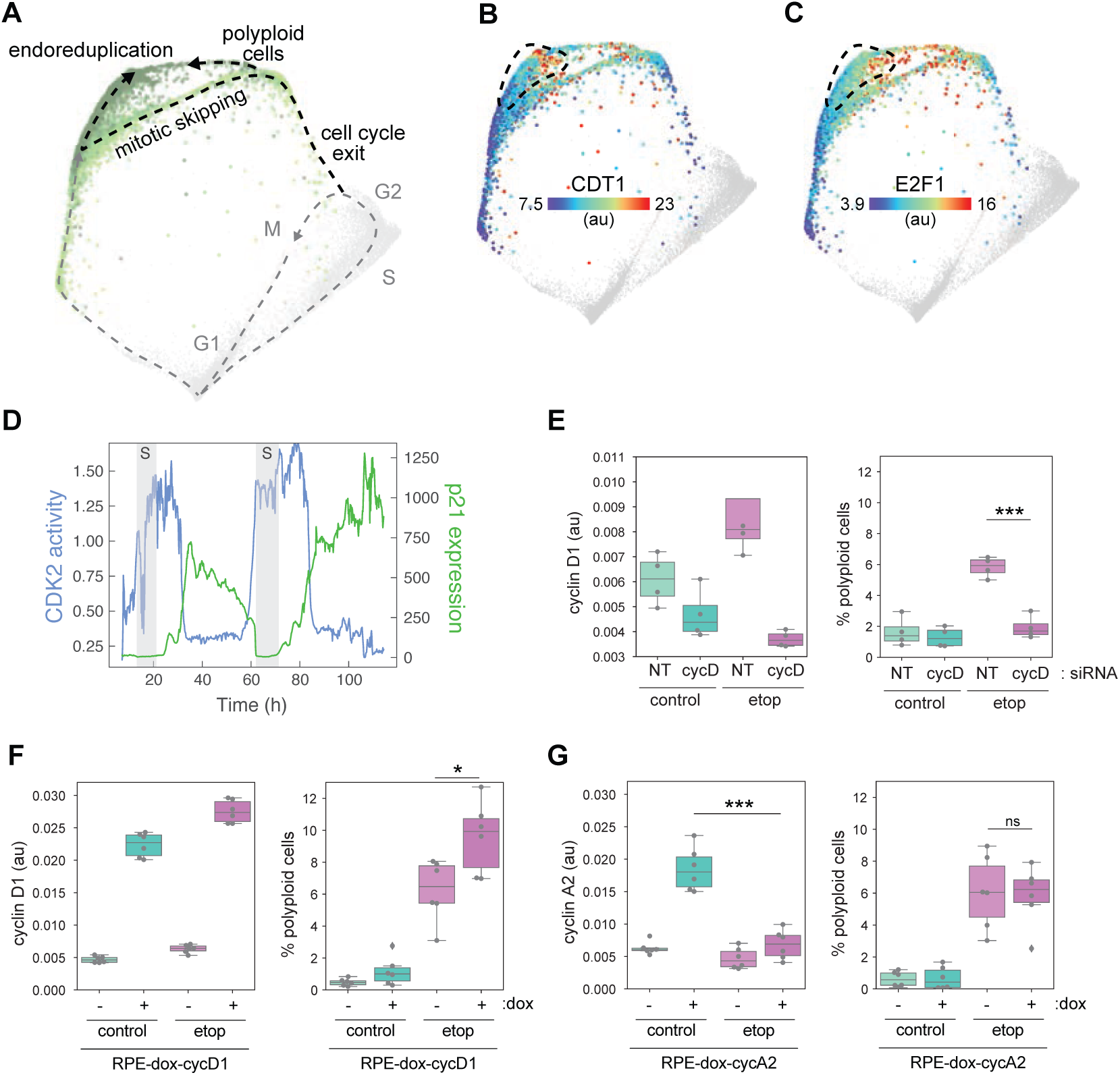
Sustained replication stress can induce polyploidy through mitotic skipping and endoreduplication. A. Arrest architecture of replicative stress. Endoreduplication and polyploid arrest trajectories are indicated. B-C. (B) CDT1 and (C) E2F1 are plotted on the map. Median nuclear values are shown. D. Representative single-cell trace demonstrating mitotic skipping and endoreduplication following etoposide treatment (1 μM). CDK2 activity (DHB-mCherry, blue), cell cycle phase (PCNA-mTq2, S phase shown as gray shaded area) and p21-YPet intensity are plotted versus time of etoposide treatment. E. Cyclin D1 abundance (left) and the proportion of polyploid cells (right), as measured by immunofluorescence and Hoechst staining, respectively, following siRNA-mediated knockdown of cyclin D in control and etoposide-treated cells. Bars represent population medians from four independent replicates (gray circles). F. Cyclin D1 abundance (left) and the proportion of polyploid cells (right), as measured by immunofluorescence and Hoechst staining, respectively, following doxycycline (dox)-induced upregulation of cyclin D1 in control and etoposide-treated cells. Bars represent population medians from six independent replicates (gray circles). G. Cyclin A abundance (left) and the proportion of polyploid cells (right), as measured by immunofluorescence and Hoechst staining, respectively, following doxycycline (dox)-induced upregulation of cyclin A2 in control and etoposide-treated cells. Bars represent population medians from six independent replicates (gray circles). Statistical significance in right panels of E-G was determined using using a two-way analysis of variance (ANOVA) with Sidak’s post hoc test (*** *p* < 0.001, * *p = 0.02*).

To test if cyclin D can drive cell cycle re-entry and endoreduplication after a G2 exit (as suggested by the structure), we treated cells with siRNA against all three cyclin D isoforms following etoposide treatment. Knockdown of cyclin D completely abolished the appearance of polyploid cells after 4 days of etoposide treatment (**Fig 6E**). Conversely, doxycycline (dox)-induced overexpression of cyclin D1 significantly increased the number of polyploid cells (**Fig 6F**), confirming its role in the generation of polyploidy. These findings indicate that increased expression of G1 cyclins in senescent cells provides a route for cells to escape from this “irreversibly” arrested state. Dox-induced expression of the S/G2 cyclin A2, on the other hand, could not increase the proportion of polyploid cells (**Fig 6G, right panel**). In fact, following etoposide treatment the dox-induced expression of cyclin A2 was significantly less than control cells following dox induction (**Fig 6G, left panel**), consistent with the reactivation of APC/C-induced degradation of cyclin A that typically occurs during G1. Thus, regardless of the phase of cell cycle exit (and DNA content), the senescent state resembles a G1-like molecular state, which narrows the mechanisms that stabilize this cell cycle arrest—as well as those that could reverse it—to G1 regulatory events.

### A map of cell cycle arrest

By projecting all three stresses onto the same structure (**Fig 7A-B**) and overlaying our inferred trajectories (**Fig 7C**), a comprehensive architecture of cell cycle arrest emerged (**Fig 7A**). The arrest architectures of replication and oxidative stress exhibited a high degree of similarity notably featuring two paths of cell cycle exit from either G1 or G2. Hypomitogenic arrest, on the other hand, induced a distinct arrest architecture with cells diverging during G2 and undergoing mitosis directly into a different state of arrest with 2C DNA content. The majority of spontaneously arrested cells (unperturbed cells with low RB phosphorylation) were observed along a trajectory towards the 2C arrest state driven by an increase in p21 (**Fig 1H**), consistent with observations that spontaneous cell cycle arrest results from low levels of replication stress (Arora *et al*, 2017).

**Fig 7:**
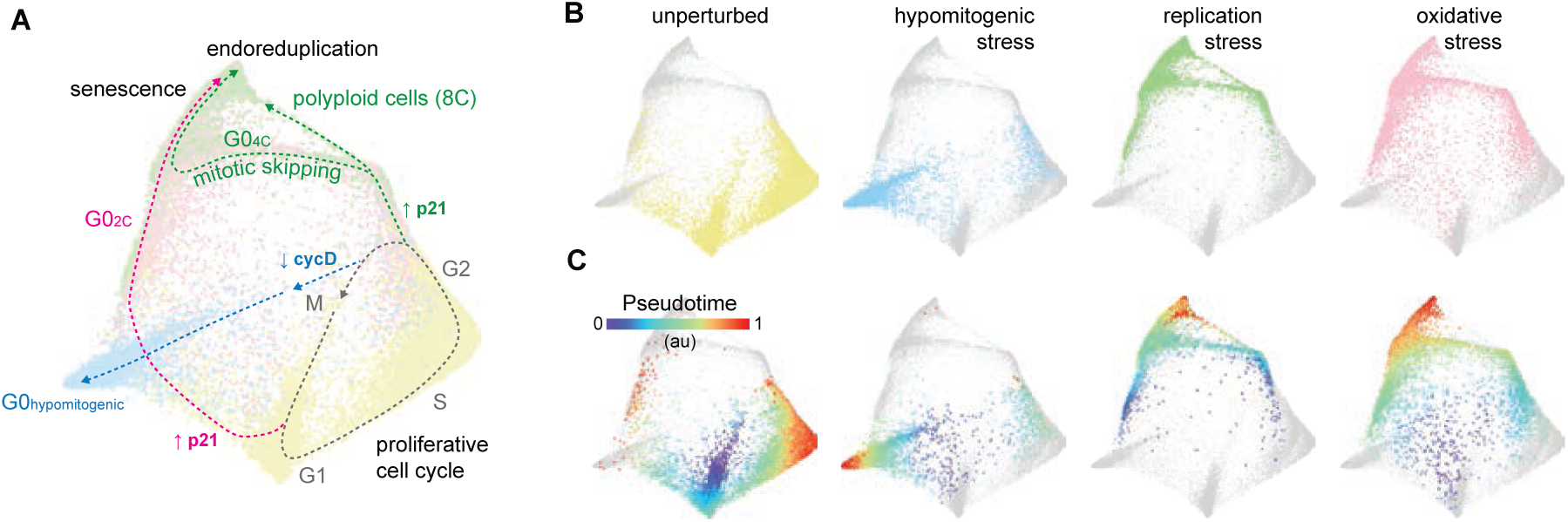
The overall architecture of cell cycle arrest. A-B. Summary map of cell cycle arrest. Unperturbed (yellow), serum-starved (blue), etoposide-(green) and H_2_O_2_-treated (pink) cells are plotted on a (A) single map. (B) Each condition is shown individually on the map (other conditions are shown in lighter gray). Inferred trajectories of the proliferative cell cycle and all arrest trajectories are shown. C. Diffusion pseudotime is plotted onto the map of each condition.

## Discussion

Here we combined hyperplexed, single-cell imaging with manifold learning to visualize the molecular architecture of cell cycle arrest and obtain a better understanding of the diversity of molecular mechanisms that govern it. In response to hypomitogenic stress, cells diverged from the proliferative trajectory in G2 lacking sufficient cyclin D1 (**Fig 2D** **and** **Fig 2G**) to sustain RB phosphorylation through mitosis as G2/M cyclins are degraded (Yang *et al*, 2017; Moser *et al*, 2018; Gookin *et al*, 2017; Stallaert *et al*, 2021). Mitogenic signaling regulates global protein translation rates throughout the cell cycle to control cyclin D abundance in mother cell G2 and influence the proliferation/arrest decision of daughter cells (Min *et al*, 2020). We found that decreased mitogenic signaling through serum starvation induced a reduction in nearly all proteins measured (including cyclin D1) as cells exited the cell cycle (**Fig 2H**), consistent with a inhibition of global protein synthesis due to a decrease in ribosomal RNA and protein synthesis (Donati *et al*, 2011) and cap-dependent translation (Liu & Qian, 2014). The abundance of p27, on the other hand, not only resisted this downregulation but was actually increased following serum starvation, likely due to the presence of an internal ribosome entry site (IRES) in the 5′-UTR of its mRNA, permitting cap-independent translation (Millard *et al*, 1997; Miskimins *et al*, 2001; Jiang *et al*, 2007), which often allows selective translation of specific proteins in conditions of cellular stress (Sonenberg & Hinnebusch, 2009). This steady increase in p27 as cells progressed further along the hypomitogenic arrest trajectory is also consistent with a molecular state that could increase the depth of cell cycle arrest (Binné *et al*, 2007; Fujimaki *et al*, 2019).

Following replication and oxidative stress, cells diverged from the proliferative trajectory at two distinct points: either (1) immediately following cell division or (2) after DNA replication (**Fig 3**). In both scenarios, cell cycle exit is driven by a DNA damage response and p21 induction (**Fig 3F**). The cell fate decision between these two trajectories in response to replication stress was predicted by the timing of the p21 induction in individual cells (**Fig 3F**), as previously shown (Barr *et al*, 2017). We found that cells that exited from G2 (along the 4C trajectory) transitioned from a G2-to-G1-like molecular state due to mitotic skipping, in which the APC/C ubiquitin ligase is activated in the absence of cell division (**Fig 4L-O**). It was previously reported these mitotic skipping mechanisms can precede senescence following a G2 cell cycle exit (Wiebusch & Hagemeier, 2010; Suzuki *et al*, 2012; Johmura *et al*, 2014; Müllers *et al*, 2014; Krenning *et al*, 2014). We directly resolved this molecular trajectory on the cell cycle map and found that it eventually converged with the arrest trajectory of cells that exited the cell cycle after cell division. Both trajectories tended toward a single, terminal senescent state with a G1-like molecular signature (**Fig 4A**).

We present several lines of evidence that senescence, under these experimental conditions, is an obligate G1-like state. First, we find that regardless of the phase of cell cycle exit following etoposide, cells accumulate over time in a single senescent state characterized by high expression of G1 effectors (**Fig 4**). In addition, we observed a similar convergence on a single senescent state following treatment with palbociclib and etoposide, which induced cell cycle exit primarily from G1 and G2, respectively (**Fig 5**). Although this state of arrest was stable under treatment, it was not completely irreversible (**Fig 6D**) and cell cycle re-entry could be induced by increased expression of the G1 effector cyclin D1 (**Fig 6E-F**). In contrast, overexpression of the S/G2 effector cyclin A2 had no effect on cell cycle re-entry (**Fig 6G**). This G1-like senescence signature with high G1 cyclins, low G2/M cyclins and 4 copies of DNA was also detected in osteosarcoma (U-2 OS) cells (**Fig EV4**).

Our results call into question the “irreversibility” of cell cycle arrest in senescence. We demonstrate that cell cycle re-entry can occur in cells that otherwise possess established hallmarks of senescence (e.g. SA-β-gal activity (Hjelmeland *et al*, 1999) and low DNA:cytoplasm ratio (Neurohr *et al*, 2019)). These re-entry events are relatively rare (5/119 cells over 4 days of sustained replication stress) but can be induced by increased expression of G1 effectors such as cyclin D1. These results are consistent with senescent cell cycle arrest being a strong attractor state (Chong *et al*, 2018; Choi *et al*, 2012), but not irreversible. Under normal physiological conditions, this senescence attractor may be sufficiently robust to maintain cell cycle arrest during normal biochemical variability. However, the supraphysiological concentrations of mitogens in which laboratory tissue cultures are grown, which stimulate the expression of cyclin D and other proliferative factors (Lukas *et al*, 1996), may allow cells to access additional states that can escape this attractor. It is possible that oncogenic transformation, which often involves the amplification of mitogenic signaling, might also increase access to these biochemical states and allow cell cycle re-entry from senescence. Furthermore, many cancer therapies are designed to induce irreversible cell cycle arrest (or death) through the generation of DNA damage, including radiotherapy and DNA-damaging agents such as etoposide. However, even a low rate of cell cycle re-entry from this arrest state could lead to tumor recurrence. Given the role that cyclin D:CDK4/6 plays in cell cycle re-entry from senescence (**Fig 6E-F**), it is conceivable that sequential treatment with a DNA-damaging agent followed by an FDA-approved CDK4/6 inhibitor such as palbociclib might help reduce tumor recurrence.

Finally, we assert that senescence is an obligate G1-like state as defined by the molecular signature of cell cycle proteins, which consequently governs the mechanisms that maintain arrest (or allow escape from it). While we also observed that many signaling and SASP markers are also similar between palbociclib- and etoposide-induced senescence (**Fig 5F-K**), it is possible that these senescent states are distinct from one another in other features not measured here, which may generate differences in other functional aspects of the senescent state (e.g. metabolism, secretory phenotype, etc). It is unclear if these other functional properties of senescent cells are also reversed when they re-enter the cell cycle.

## Materials and Methods

### Cell lines and culture

Retinal pigment epithelial cells (hTERT RPE-1, ATCC, CRL-4000) were used for all experiments except time-lapse imaging. The RPE-PCNA-mTq2/p21-YPet/DHB-mCherry (Stallaert *et al*, 2021) and RPE-p21-mTq2/cycD1-mVenus/DHB-mCherry/H2B-mIFP (Zerjatke *et al*, 2017) cell lines were previously described. The RPE-mVenus-p27K-cell line was created using the pMXs-IP-mVenus-p27K− plasmid and was described in (Oki *et al*, 2014). It encodes retrovirus packaging sequences to package a non-CDK binding p27 mutant fused to mVenus for expression under control of a constitutive heterologous promoter. As reported, mouse p27 was amplified from mouse bone marrow cDNA using primers with 5′-XhoI and 3′-NotI sites by PCR. The p27K− were generated by PCR-based mutagenesis. The constructs were subcloned into a pMXs-internal ribosome entry site (IRES)-Puro vector (pMXs-IP). To make the RPE1-hTert line containing PCNA-mTurq2, p27-mVenus, and DHB-mCherry, the plasmids were transfected into HEK293T with pCI-GPZ or ΔNRF and Vesicular stomatitis virus G packaging plasmids with 50 μg/ml polyethylenimine (Aldrich Chemistry). Viral supernatants were used to transduce RPE1-hTert cells in the presence of 8 µg/ml polybrene (Millipore). A clonal cell line was picked based on fluorescence of all three biosensors.

All cells were grown at 37°C and 5% CO2 in DMEM (Gibco, 11995-065) with 10% fetal bovine serum (FBS; Sigma, TMS-013-B), 2 mM L-glutamine (ThermoFisher Scientific, 25030081) and penicillin/streptomycin (P/S; ThermoFisher Scientific, 15140148). FluoroBrite™ DMEM (Gibco, A18967-01) supplemented with 10% FBS and 2 mM L-glutamine was for time-lapse imaging. All cell lines were authenticated by STR profiling (ATCC) and confirmed to be mycoplasma free. Where indicated, cells were treated with etoposide (MedChemExpress, HY-13629), H_2_O_2_ (Millipore Sigma, H1009) or palbociclib (Selleckchem, S1116).

### Antibodies

All antibodies used in this study were chosen using BenchSci (http://app.benchsci.com) to identify high quality, previously published/validated primary antibodies and are listed in Table EV1.

### Time-lapse imaging

Time-lapse imaging was performed as previously described (Stallaert *et al*, 2021) using a Nikon Ti Eclipse inverted microscope equipped with a Nikon Plan Apochromat Lambda 40x objective (NA=0.95) and an Andor Zyla 4.2 sCMOS detector, using Nikon Perfect Focus System (PFS). A climate-controlled enclosure (Okolabs) was used to maintain constant temperature (37°C) and atmosphere (5% CO_2_). Combinations of the following filter sets (Chroma) were used as required (excitation; beam splitter; emission filter): CFP (425-445/455/465-495nm), YFP (490-510/515/520-550nm), mCherry(540-580/585/593-668) and Cy5(590-650/660/663-738nm).

RPE-PCNA-mTq2/p21-YPet/DHB-mCherry and RPE-p21-mTq2/cycD1-mVenus/DHB-mCherry/ H2B-mIFP cells were imaged every 10 min and RPE-mVenus-p27K-cells were imaged every 15 minutes. Stitched 4×4 images were acquired and field illumination correction was performed before stitching. Image analysis and post-processing was performed using NIS-Elements AR software with General Analysis 3. CDK2 activity was calculated as the ratio of background corrected cytoplasmic to nuclear intensity of DHB-mCherry. The cytoplasm signal was quantified in a 15-pixel ring outside the segmented nucleus, with a 2-pixel gap between the nucleus and the ring. p21-YPet or cyclinD1-mVenus intensity was calculated as the background corrected median nuclear intensity. The appearance/disappearance of nuclear PCNA foci was used to manually annotate cell cycle phase transitions.

Image analysis was performed using Python (3.7.10) with Cellpose (v. 0.6.5) (Stringer *et al*, 2021) segmentation algorithm, Scikit-image image processing library (v.0.18.2) (van der Walt *et al*, 2014) and BayesianTracker linking algorithm (v. 0.4.1) (Ulicna *et al*, 2021). Errors in segmentation and tracking were corrected manually using the napari graphical interface (v. 0.4.10) (Sofroniew *et al*, 2022).

### Iterative indirect immunofluorescence imaging (4i)

Sample preparation was performed as previously described (Stallaert *et al*, 2021). Stitched 8×8 images were acquired for each condition using the Nikon Ti Eclipse microscope described above with the following filter cubes (Chroma): DAPI(383-408/425/435-485nm), GFP(450-490/495/500-550nm), Cy3(530-560/570/573-648nm) and Cy5(590-650/660/663-738nm). Images from successive rounds were aligned in Python (v3.7.1) using the StackReg library (Thévenaz *et al*, 1998) and corrected using manually-selected fiduciary points if necessary. Segmentation and feature extraction were performed in CellProfiler (v3.1.8) (Stirling *et al*, 2021) using standard modules.

### Data integration

The following normalization was used to integrate the independent datasets containing 4i measurements of (1) unperturbed (control 1), serum-starved, etoposide- and H_2_O_2_-treated cells and (2) unperturbed (control 2) and palbociclib-treated cells. For each of the datasets, first the control data was z-normalized. Each treatment condition was subsequently normalized using the mean and standard deviation of its matched control, to preserve relative fold-changes induced by treatment. Bimodal features used for subsequent embedding were peak normalized (2C and 4C peaks in DNA content normalized to 2 and 4, phospho-RB and phospho/total RB hypo- and hyperphosphorylation peaks normalized to 0 and 1).

### Feature selection and manifold learning

We previously trained random forest models on ground truth annotations of cell cycle phase and age to identify an optimized feature set that resolves the cell cycle manifold in RPE cells (Stallaert *et al*, 2021). This validated feature set was used to inform feature selection in the current study (**Dataset EV1**). Manifold learning was performed using Potential of Heat-diffusion for Affinity-based Transition Embedding (PHATE) (Moon *et al*, 2019) using the feature set described above as input variables. PHATE was run on z-normalized features with the following parameter sets for cell cycle maps: (Fig 1B: k-nearest neighbor (knn)=150, t=20, gamma=1; Fig 2A: knn=15, t=26, gamma=1, Fig 3A: knn=75, t=19, gamma=1, Fig 5A: knn=50, t=19, gamma=1, Fig 7A: knn=150, t=10, gamma=0.25, Fig EV5A: knn=150, t=10, gamma=0.25, Fig EV3A: knn=100, t=30, gamma=1).

### Data Visualization

Python (v3.7.1) and Jupyter Notebooks (v6.1.4) were used for data visualization using matplotlib (v3.3.2), seaborn (v0.11.0) and scanpy (v1.6) (Wolf *et al*, 2018) libraries, as well as GraphPad Prism (v8).

### siRNA

RPE cells were treated with DMSO or etoposide (1 μM) for 24h then transfected with non-targeting (Dharmacon, D-001810-10-0) or cyclin D1/D2/D3 (SMARTPools L-003210-00-0005/L-003211-00-0005/L-003212-00-0005) siRNA pools using the DharmaFECT 1 transfection reagent (T-2001-01) as per manufacturer’s protocol and incubated for 3 days prior to fixation. Immunofluorescence was performed as per the 4i protocol described above.

## Acknowledgments

We would like to thank Dr. Sam Wolff (UNC Chapel Hill) for support with imaging. The RPE-p21-mTq2/cycD1-mVenus/DHB-mCherry/H2B-mIFP was a generous gift from Dr. Jorg Mansfield (Institute for Cancer Research, London, England) and Dr. Sabrina Spencer (University of Colorado Boulder). The pMXs-IP-mVenus-p27K− plasmid used to make the RPE-mVenus-p27K-cell line was a gift from T. Kitamura (University of Tokyo, Minato-ku, Tokyo).

This work was supported by NSF CAREER Award 1845796 (JEP); NIH grants R01-GM138834 (JEP), R01-GM083024 (JGC), R01-GM102413 (JGC), and R35-GM141833 (JGC); the Chan Zuckerberg Initiative DAF (an advised fund of Silicon Valley Community Foundation 2020-225716) (KMK); JCL was supported by an HHMI Gilliam Fellowship for Advanced Study (GT10886); and NIH/NIGMS awards to UNC: R25-GM055336 and T32-GM007040.

## Author contributions

WS and JEP conceived of the project. WS, KMK, JGC and JEP designed the experiments. WS, HKS, SRT and CLY performed the 4i experiments. WS, MSJ and SRT performed the time-lapse imaging. WS, KMK, JH, MSJ and CDT performed image analysis. MSJ and JCL created cell lines. WS wrote the manuscript with the help of all authors.

## Data and materials availability

Single cell datasets are available at doi:10.5281/zenodo.6394367.

---

**Fig EV1:**
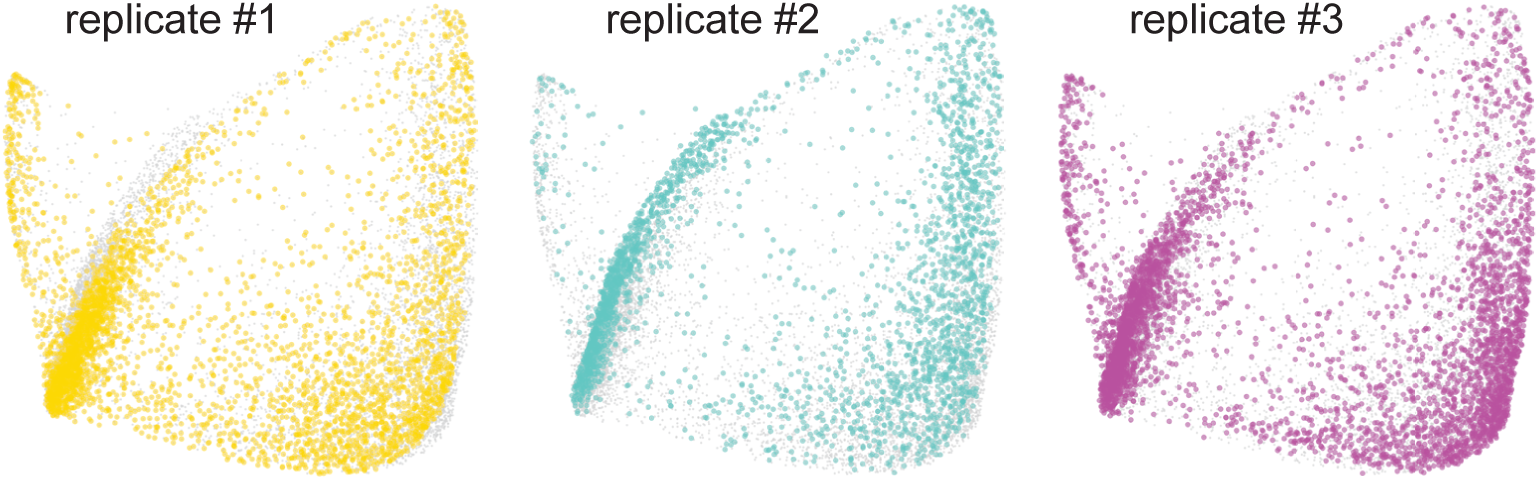
Replicates of the unperturbed cell cycle map.

**Fig EV2:**
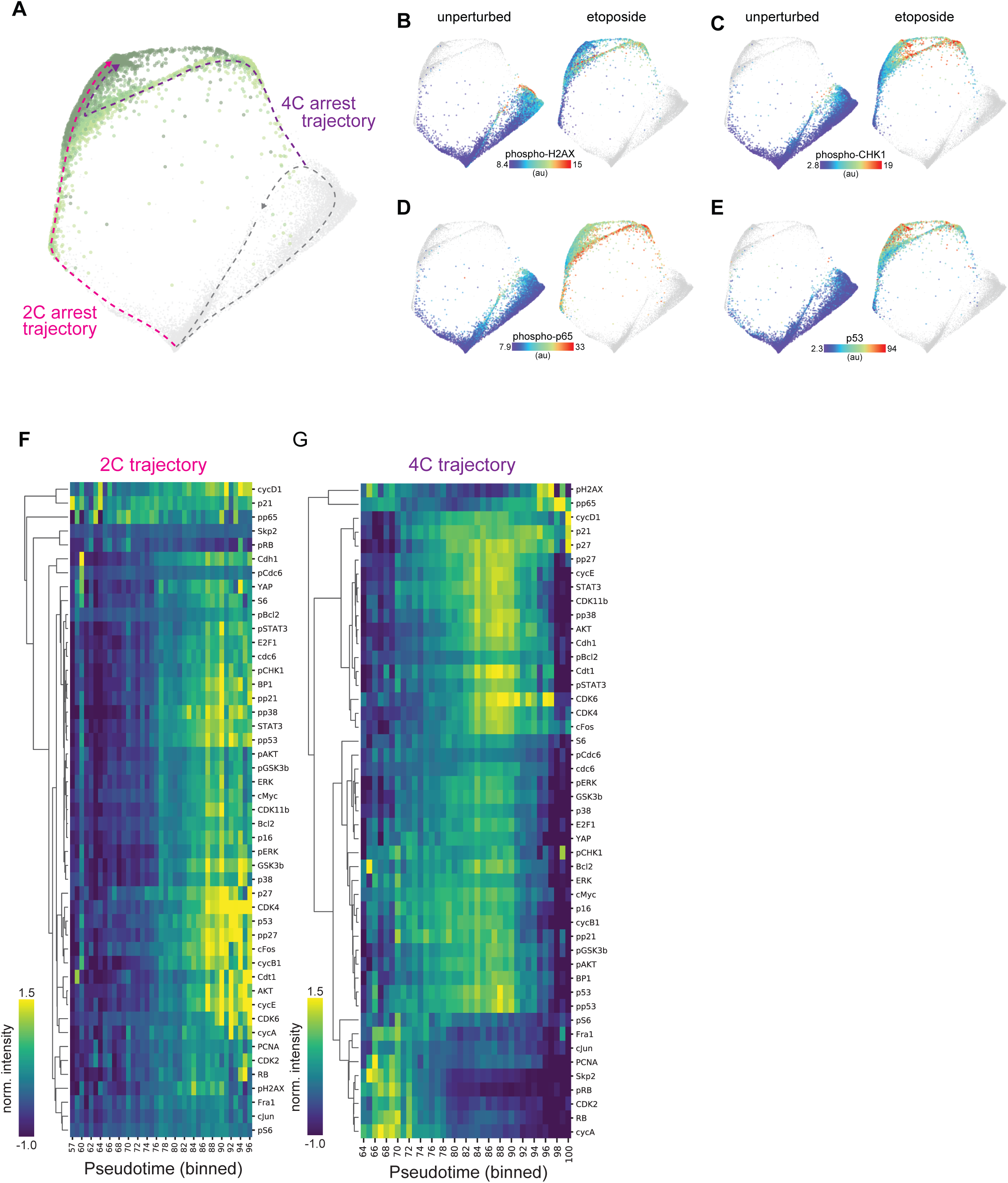
Arrest trajectories following replication stress. A. Cell cycle map arrest of unperturbed (gray) and etoposide-treated cells (1 μM; 1d: light green, 2d: green, 3d: dark green, 4d: darker green). The unperturbed cell cycle (dotted gray line) and two arrest trajectories (into 2C and 4C, pink and purple, respectively) are indicated on the map. B-E. (B) Phospho-H2AX, (C) phospho-CHK1, (D) phospho-p65 and (E) p53 of unperturbed (left panels) or etoposide-treated cells (right panels) are plotted on the arrest architecture. Median nuclear values are shown. C. Heatmap of feature intensity along the 2C (left) and 4C (right) arrest trajectories. Features were ordered by hierarchical clustering according to their dynamics along each arrest trajectory. Diffusion pseudotime values were binned and pseudotime values with <15 cells were excluded from the visualization.

**Fig EV3:**
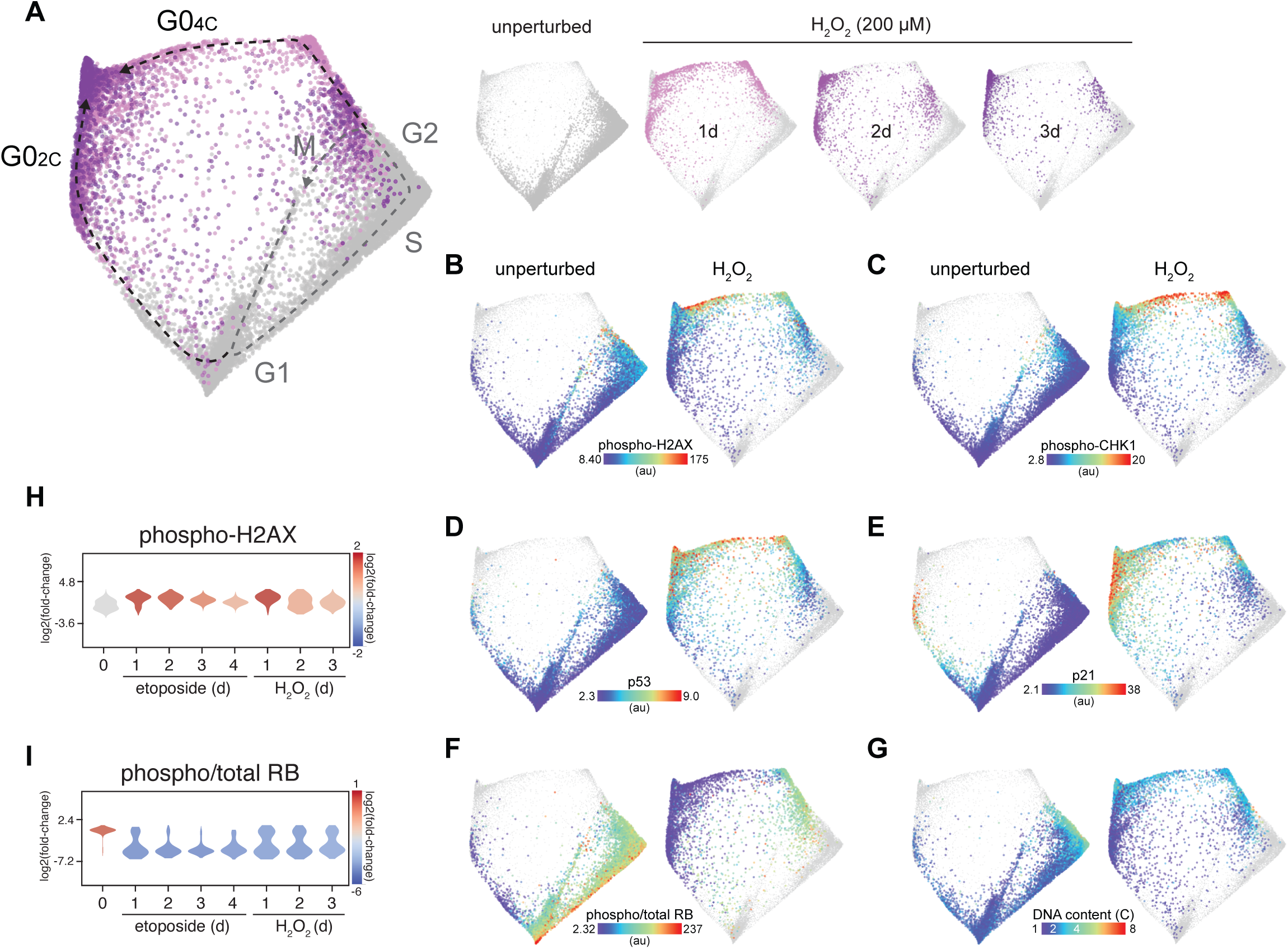
The arrest architecture of oxidative stress. B. Unified cell cycle map arrest of unperturbed (gray) and H_2_O_2_-treated cells (200 μM; 1d: light purple, 2d: purple, 3d: dark purple – see inset, *N* = 5015 cells). The unperturbed cell cycle (dotted gray line) and two arrest trajectories (into G0_2C_ and G0_4C_; black dotted lines) are indicated on the map. Inset: Each condition is shown individually on the map (other conditions are shown in lighter gray). B-G. (B) Phospho-H2AX, (C) phospho-CHK1, (D) p53, (E) p21, (F) phospho/total RB and (G) DNA content of unperturbed (left panels) or H_2_O_2_-treated cells (right panels) are plotted on the arrest architecture. Median nuclear values are shown. H-I. Distribution of (H) phospho-H2AX and (I) phospho/total RB in individual cells following etoposide (1 μM) or H_2_O_2_ treatment (200 μM).

**Fig EV4:**
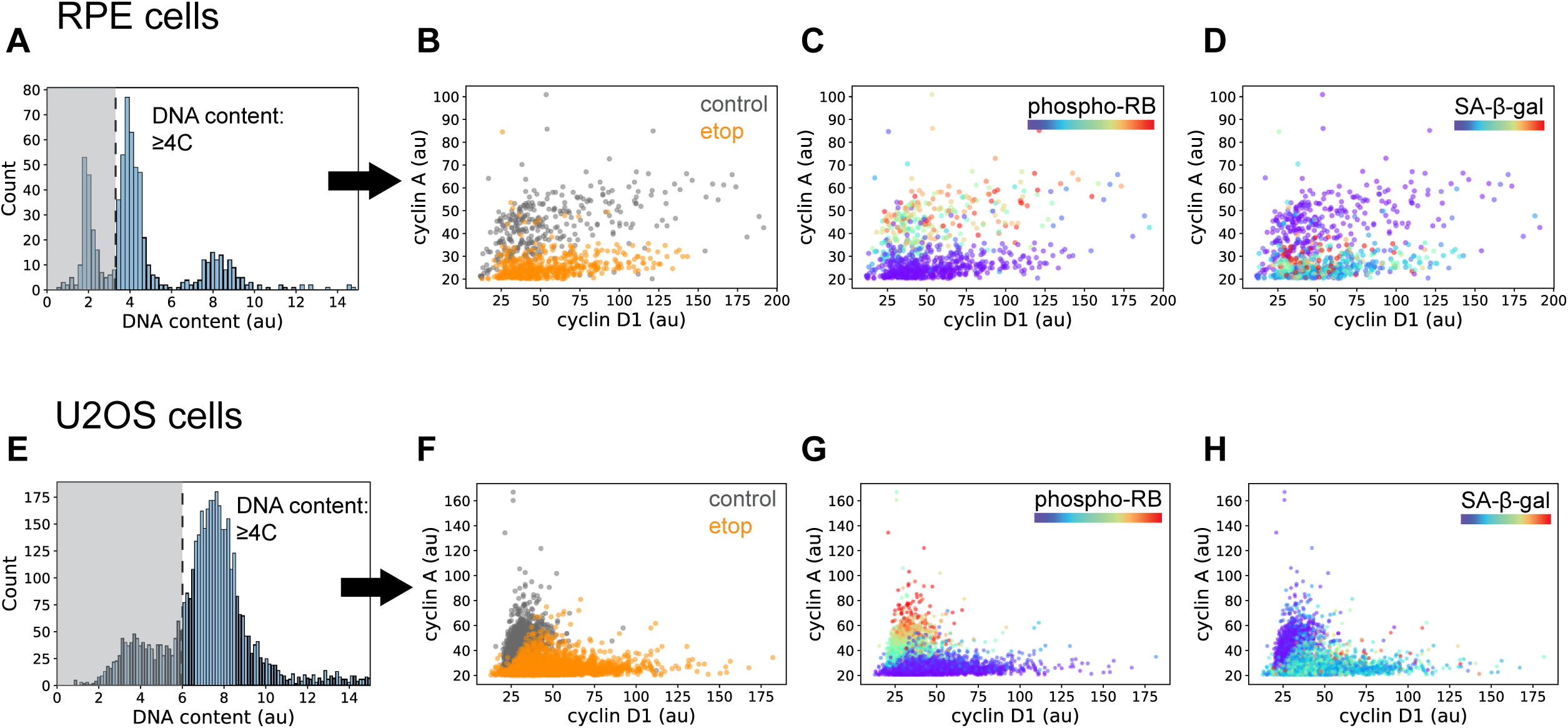
Replication stress induces mitotic skipping in RPE and U-2 OS cells. A-D. (A) RPE cells with ≥4C DNA content were selected using the intensity of nuclear Hoechst staining. Cyclin A versus cyclin D1 intensity was plotted for each cell and overlaid with (B) condition labels (control vs. etoposide), (C) phospho-RB intensity and (D) senescence-associated β-galactosidase (SA-βgal) activity. E-H. The same analysis as above was performed on U-2 OS cells.

**Fig EV5:**
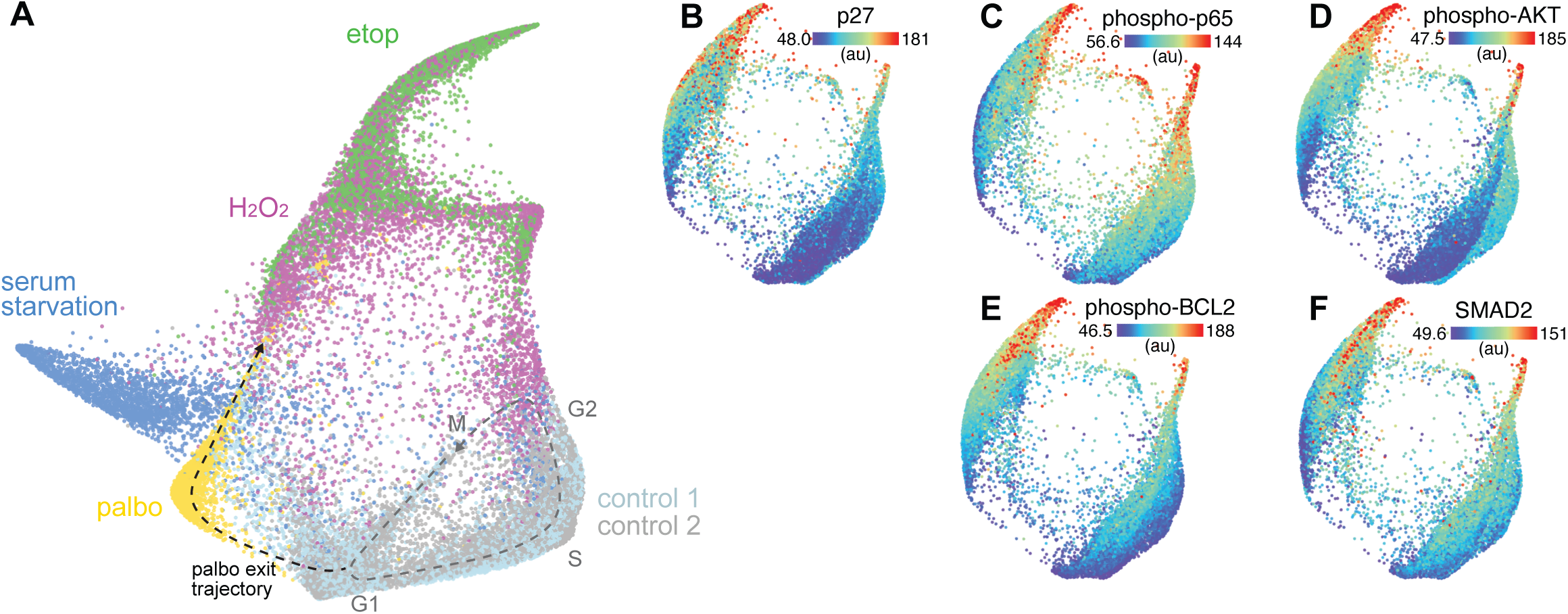
The arrest architecture of palbociclib-induced arrest. A. Unified cell cycle map of unperturbed (control 2, gray) and palbociclib-treated cells (gold) from a separate experiment, plotted with unperturbed (control 1: light blue, from original experiment), serum-starved (blue) etoposide-(etop, green) and H_2_O_2_-treated (magenta) cells from initial experiment. Data integration is described in Materials and Methods. B-F. (B) p27, (C) phospho-p65, (D) phospho-AKT, (E) phospho-BCL2 and (F) SMAD2 are plotted on the map. Median nuclear values are shown.

**Table EV1.** Antibodies

**Primary Antibodies**
| Target | Host species | Dilution | Supplier | Catalog# | Published citations | Published figures |
| --- | --- | --- | --- | --- | --- | --- |
| 53BP1 | Rb | 1:1000 | Santa Cruz | sc-22760 | 114 | 207 |
| Akt | Ms | 1:100 | CST | 2920 | 397 | 649 |
| Bcl-2 | Rb | 1:200 | CST | 2827 | 38 | 60 |
| c-Fos | Rb | 1:200 | CST | 2250 | 81 | 124 |
| c-Jun | Rb | 1:400 | CST | 9165 | 356 | 620 |
| c-Myc | Rb | 1:500 | CST | 5605 | 402 |  |
| cdc6 | Ms | 1:100 | Santa-Cruz | sc-9964 | 33 | 88 |
| Cdh1 | Ms | 1:100 | Santa Cruz | sc-56312 | 6 | 8 |
| CDK2 | Gt | 1:500 | R&D | AF4654 | 2 | 2 |
| CDK4 | Rb | 1:400 | abcam | ab108355 | 3 | 3 |
| CDK6 | Rb | 1:250 | abcam | ab124821 | 19 | 30 |
| CDK11b | Rb | 1:100 | Abcam | ab198813 | 1 | 2 |
| Cdt1 | Rb | 1:200 | CST | 8064 | 9 | 28 |
| Cyclin A | Ms | 1:50 | Santa Cruz | sc-271682 | 13 | 18 |
| cyclin B1 | Gt | 1:100 | R&D | AF6000 | 41 | 52 |
| cyclin D1 | Ms | 1:100 | Santa Cruz | sc-20044 | 115 | 190 |
| cyclin E | Ms | 1:50 | Santa Cruz | sc-247 | 47 | 66 |
| E2F1 | Ms | 1:100 | Santa Cruz | sc-251 | 146 | 273 |
| ERK | Ms | 1:100 | CST | 4696 | 132 | 181 |
| Fra1 | Ms | 1:100 | Santa Cruz | sc-28310 | 23 | 70 |
| GSK3b | Ms | 1:100 | CST | 9832 | 64 | 113 |
| p16 | Rb | 1:400 | abcam | ab108349 | 40 | 62 |
| p21 | Gt | 1:200 | R&D | AF1047 | 7 | 18 |
| p27 | Rb | 1:800 | CST | 3686 | 58 | 85 |
| p38 | Rb | 1:100 | CST | 8690 | 426 | 573 |
| p53 | Ms | 1:400 | CST | 48818 | 12 | 23 |
| PCNA | Ms | 1:500 | CST | 2586 | 193 | 286 |
| phospho-Akt (Ser473) | Rb | 1:200 | CST | 4060 | 2700 | 4300 |
| phospho-Bcl2 | Ms | 1:200 | CST | 15071 | 159 | 220 |
| phospho-cdc6 (Ser54) | Rb | 1:200 | abcam | ab75809 | 4 | 13 |
| phospho-CHK1 (S317) | Rb | 1:800 | CST | 12302 | 17 | 30 |
| phospho-ERK (T202/Y204) | Rb | 1:200 | CST | 4370 | 1600 | 2300 |
| phospho-H2AX (S139) | Ms | 1:200 | CST | 80312 | 3 | 3 |
| phospho-GSK3b (S9) | Rb | 1:200 | CST | 5558 | 188 | 271 |
| phospho-p21 (T145) | Rb | 1:1000 | abcam | ab47300 | 2 | 4 |
| phospho-p27 (T157) | Rb | 1:60 | abcam | ab85047 | 5 | 5 |
| phospho-p38 (T180/Y182) | Ms | 1:200 | CST | 9216 | 158 | 250 |
| phospho-p53 (S15) | Ms | 1:400 | CST | 9286 | 87 | 145 |
| phospho-p65 (S536) | Ms | 1:100 | CST | 3036 | 32 | 44 |
| phospho-RB | Rb | 1:1000 | CST | 8516 | 48 | 70 |
| phospho-S6 (S235/236) | Rb | 1:200 | CST | 4858 | 156 | 250 |
| phospho-STAT3 (Y705) | Rb | 1:100 | CST | 9145 | 1000 | 1700 |
| RB | Ms | 1:500 | CST | 9309 | 123 | 180 |
| Rsk1 | Ms | 1:50 | R&D | MAB992 |  |  |
| S6 | Ms | 1:100 | CST | 2317 | 107 | 152 |
| Skp2 | Rb | 1:800 | CST | 2652 | 20 | 47 |
| SMAD2 | Ms | 1:200 | abcam | ab119907 | 3 | 8 |
| STAT3 | Ms | 1:500 | CST | 9139 | 551 | 1000 |
| YAP | Ms | 1:50 | Santa Cruz | sc-101199 | 55 | 129 |

**Secondary Antibodies**
| Host/target species | Fluorophore | Dilution | Supplier | Catalog# |
| --- | --- | --- | --- | --- |
| Donkey anti-rabbit | AlexaFluor Plus 488 | 1:500 | ThermoFisher | A32790 |
| Donkey anti-mouse | AlexaFluor Plus 555 | 1:500 | ThermoFisher | A32773 |
| Donkey anti-rabbit | AlexaFluor Plus 647 | 1:500 | ThermoFisher | A32773 |
| Donkey anti-goat | AlexaFluor Plus 647 | 1:500 | ThermoFisher | A32758 |

**Dataset EV1:** PHATE input features

|  |  |
| --- | --- |
| AreaShape_Area_cell | Int_Med_cycD1_nuc |
| AreaShape_Area_nuc | Int_Med_cycE_nuc |
| Int_Intg_DNA_nuc | Int_Med_p16_nuc |
| Int_Med_BP1_nuc | Int_Med_p21_nuc |
| Int_Med_CDK11b_nuc | Int_Med_p27_nuc |
| Int_Med_CDK2_nuc | Int_Med_p38_nuc |
| Int_Med_CDK4_nuc | Int_Med_p53_nuc |
| Int_Med_CDK6_nuc | Int_Med_pCHK1_nuc |
| Int_Med_Cdh1_nuc | Int_Med_pCdc6_nuc |
| Int_Med_Cdt1_nuc | Int_Med_pH2AX_nuc |
| Int_Med_DNA_nuc | Int_Med_pRB_nuc |
| Int_Med_E2F1_nuc | Int_Med_pS6_nuc |
| Int_Med_Fra1_nuc | Int_Med_pp21_nuc |
| Int_Med_RB_nuc | Int_Med_pp27_nuc |
| Int_Med_S6_nuc | Int_Med_pp38_nuc |
| Int_Med_Skp2_nuc | Int_Med_pp53_nuc |
| Int_Med_YAP_nuc | Int_Med_PCNA_nuc |
| Int_Med_cFos_nuc | Int_Std_PCNA_nuc |
| Int_Med_cJun_nuc | Int_pRB_over_RB_nuc |
| Int_Med_cMyc_nuc | Int_pS6_over_S6_nuc |
| Int_Med_cdc6_nuc | Int_pp21_over_p21_nuc |
| Int_Med_cycA_nuc | Int_pp27_over_p27_nuc |
| Int_Med_cycB1_nuc | Int_pp53_over_p53_nuc |
|  | Int_pp38_over_p38_nuc |

